# The Instability of Functional Connectomes Across the First Year of Life

**DOI:** 10.1101/2021.04.14.439877

**Authors:** Alexander J. Dufford, Stephanie Noble, Siyuan Gao, Dustin Scheinost

## Abstract

The uniqueness and stability of the adolescent and adult functional connectome has been demonstrated to be high (80–95% identification) using connectome-based identification (ID) or “fingerprinting”. However, it is unclear to what extent individuals exhibit similar distinctiveness and stability in infancy, a developmental period of rapid and unparalleled brain development. In this study, we examined connectome-based ID rates within and across the first year of life using a longitudinal infant dataset at 1 month and 9 months of age. We also calculated the test–retest reliability of individual connections across the first year of life using the intraclass correlation coefficient (ICC). Overall, we found substantially lower infant ID rates than have been reported in adult and adolescent populations. Within-session ID rates were moderate and significant (ID = 48.94–70.83%). Between-session ID rates were very low and not significant, with task-to-task connectomes resulting in the highest between-session ID rate (ID = 26.6%). Similarly, average edge-level test-retest reliability was higher within-session than between-session (mean within-session ICC = 0.17, mean between-session ICC = 0.10). These findings suggest a lack of uniqueness and stability in functional connectomes across the first year of life consistent with the unparalleled changes in brain functional organization during this critical period.

## Introduction

The first year of life is marked by rapid and unparalleled rates of brain development (Gao, Lin, Grewen, & Gilmore, 2017; Holland et al., 2014). Across the first year of life, functional networks mature towards ‘adult-like’ connectivity patterns from primary and secondary visual networks, to higher order networks such as the dorsal attention network and default mode network (Gao, Alcauter, Elton, et al., 2015; Gao, Alcauter, Smith, Gilmore, & Lin, 2015). While functional connectivity is rapidly developing in infancy, it has been found to be both unique and stable in adults (Finn et al., 2015; Horien et al., 2018) and adolescents (Horien, Shen, Scheinost, & Constable, 2019; Jalbrzikowski et al., 2019; Kaufmann et al., 2017). This uniqueness and stability in adulthood has been demonstrated using connectome-based identification (ID) or “fingerprinting”, a procedure which uses functional connectomes to identify an individual from a pool of other individuals based upon the similarity of connectomes from separate sessions. Examining the uniqueness and stability of the infant functional connectome has the potential to provide insight into how the functional connectome develops its individual variability (Finn et al., 2015). Given the rapid and dynamic changes to the brain’s functional architecture during this period (Gao, Alcauter, Smith, et al., 2015; Gao et al., 2017; Zhang, Shen, & Lin, 2019), understanding the development of its individual variability may be useful for understanding both typical and atypical neurodevelopment. Studies using this connectome-based ID procedure have found ID rates between 80–95% in adults and adolescents (Finn et al., 2015; Horien et al., 2018; Horien et al., 2019; Jalbrzikowski et al., 2019; Kaufmann et al., 2017). However, it is unclear if high ID rates are achievable in infancy when functional connectivity is undergoing extensive development.

A complementary approach is to examine the stability of individual connections, or edges. In adults, edge-level test-retest reliability via the intraclass correlation coefficient (ICC) is typically low despite high ID rates using the full connectome (Noble et al., 2017). A recent study of test–retest reliability in infants provides evidence that edge intraclass correlation for infants is low (Wang et al., 2020). However, this study only included resting-state data, tested within-session test–retest reliability and did not compare edge-level results with connectome-based ID, which measures similarity using the full pattern of the connectome. In this study, we expand upon these findings and examine test–retest reliability for both resting-state and task-based data as well as between-session test–retest reliability.

We examined these questions using data from infants that have both resting-state and task data (native language task) at 1–2 months and at 9–10 months. We hypothesized between-session connectome-based ID with functional connectomes calculated from resting-state data would be poor (i.e., ID rates below 50%); further, we hypothesized between-session ID based upon task data would be fair (60–80%) but not as high as in adult studies (90%). We expected that within-session ID would be higher than between session and overall higher in the task-based data. Regarding the test–retest reliability of the functional connectivity data, we hypothesized that edge intraclass correlation coefficients would be poor for both between-session resting-state and task-based data connectomes with task-based data being slightly higher. We predicted edge intraclass correlation coefficients would be higher for both within-session resting-state and task-based data, with task-based data being higher. Lastly, studies in adults and adolescence have found frontoparietal connections contribute most to successful ID. While these connections are immature in infancy, we expected that the immature connectivity of frontoparietal connectivity will still contribute most to successful ID with more mature patterns of connectivity being associated with higher success of ID.

## 1. Methods and Materials

### 1.1. Infant dataset (NDAR) and participants

Infant data for this study were acquired from the National Database for Autism Research (NDAR). The dataset identifier is NDARCOL0002026 (Susan Bookheimer). This study used longitudinal neuroimaging in infancy to examine early markers of Autism Spectrum Disorder (ASD). Infants were recruited in two categories: Low Risk (LR) and High Risk (HR). HR was defined as the infant having at least one older sibling with a confirmed diagnosis of ASD. Infants were categorized as HR if they had no family history of ASD defined as no first- or second-degree relatives with ASD or another neurodevelopmental disorder. Participant exclusionary criteria included: genetic or neurological condition, perinatal condition impacting development, visual, hearing, or motor impairment, and MRI contraindication. Infants were full-term and had births without complications and normal birth weight (>3000 g). At Session 1 (1.5 months), 74 infants were scanned (41 HR, 33 LR); however, 4 HR and 1 LR did not complete the scan. At Session 2 (9 months) 78 infants were scanned (48 HR, 30 LR); however, 7 HR and 5 LR infants did not complete the scan. Infants at Session 1 and 2 were scanned during rest (naturally sleep with no task) and task (native language task); however, the infants that completed these sessions successfully and had usable data varied between scan type (rest or task) and session. Two infants (HR) were removed from the analysis as testing with the ADOS indicated moderate to severe concern for ASD. Thus, data from n = 55 infants were used for subsequent analyses (n = 22 or 40% female, n = 28 or 50.91% high risk).”

For the between-session analyses (connectome-based ID and test–retest reliability) there were n = 27 for Rest1–Rest2 (had usable data at rest session 1 and rest session 2) and n = 15 for Task1–Task2. For Rest1–Task2 there were n = 24 with usable data, and n = 17 for Task1–Rest2. For the within-session analyses, there were n = 24 for Rest1–Task1 and n = 47 for Rest2–Task2. This study was approved by the Institutional Review Board of the Yale School of Medicine. The data used for the study is available at https://nda.nih.gov/. In post-analysis, we examined if successful ID varied by ASD risk status (low risk versus high risk) or infant sex (male versus female) using two Pearson’s chi-square test with Yate’s correction for continuity. The Yate’s correction is suggested when analyzing a 2×2 contingency table if one of cell frequencies is below 10 (Camilli & Hopkins, 1978).

### 1.2. Infant MRI acquisition

Infant MRI data were acquired using a 3T Siemens Tim Trio and 3T Siemens Prisma; data were acquired using a 12-channel head coil for the Tim Trio scanner, and a 32-channel head coil for the Prisma scanner. High resolution T2-weighted echo planar structural images were acquired (TR = 5000 ms, TE = 28 ms, FOV = 192, 34 slices, 128 x 128 matrix, 1.5 mm in-plane resolution, 4mm-thick axial slices for the Tim Trio, all identical parameters for the Prisma except for 33 slices were acquired and TE = 45 ms). Resting-state acquisitions were 8 minutes in duration using a T2*-weighted echo planar imaging sequence (TR = 2000 ms, TE = 28 ms, 64 x 64 matrix, FOV = 192 mm, 34 slices, 3 mm in-plane resolution, 4 mm-thick axial slices). Native language task data were acquired with a T2*-weighted echo planar imaging sequence (TR = 3000 ms, TE = 28 ms, 56 x 56 matrix, FOV = 192 mm, 34 slices, 4 mm thick for the Tim Trio, and all parameters were identical for the Prisma except for 33 slices were acquired) 7.2 minutes in duration (144 volumes).

Resting-state and native language task data were collected from infants during their natural sleep. Participants put their infant to sleep during their regular bedtime. Once asleep, the swaddled infants were transferred into the scanner. Infants were fit with earplugs and MiniMuffs (Natus Medical Inc., San Carlos, California) for hearing protection as well as headphones to convey the auditory stimuli during the native language task. Infants were laid in a custom-made bed which could fit inside the head coil and was secured to the scanner bed with Velcro. A weighted blanked and foam pads were used to minimize infant head motion. A staff member of the study remained inside the scanner room to monitor large infant motion, waking, or signal of distress i.e. crying. For the native language task, infants heard speech stimuli from different female native speakers of English and Japanese. Infants heard 7 segments in each language that were matched on duration, intensity, peak amplitude, pitch, and pitch range. The native language task used a traditional block design with alternating streams of English and Japanese speech (18 seconds in duration) interleaved with blocks of silence (12 seconds in duration). Infants heard the speech through the MRI-compatible headphones.

### 1.3. Infant functional connectivity processing

Resting-state and native language task data underwent motion correction using SPM8 (http://www.fil.ion.ucl.ac.uk/spm/). Images were iteratively smoothed using AFNI’s (http://afni.nimh.nih.gov/afni/) 3dBlurToFWHM to reach a smoothness of approximately 8 mm full-width half maximum (FWHM). Linear and quadratic drifts, mean cerebrospinal fluid signal, mean white matter signal, and mean gray matter signal were regressed from the data. Additionally, a 24-parameter motion model was regressed from the data. The 24-parameter motion model includes six rigid-body motion parameters, six temporal derivatives, and these terms squared. The functional data was temporally smoothed with a Gaussian filter with an approximate cutoff frequency = 0.12 Hz. Gray matter, white matter, and cerebrospinal fluid masks were defined on the reference brain and a dilate gray matter mask was applied to include on voxels within gray matter for further calculations. As motion has been shown to be an important confound in functional connectivity studies, the frame-to-frame displacement averaged across the functional volumes was calculated for each session of restingstate and the native language task. Infants were excluded from the analysis if their average frame-to-frame motion was greater than 0.15 mm for one of the scan types for that ID calculation. For Rest1–Task1, 15 infants were excluded due to motion, 4 for Rest2–Task2, 1 for Rest1–Rest2, 3 for Rest1–Task2, and 8 for Task1–Rest2.

### 1.4. Infant connectivity matrices

Connectivity matrices were calculated using an infant-specific parcellation which consisted of 95 nodes providing whole brain coverage (Scheinost et al., 2016). First, the parcellation was defined in the template space and then transformed back into each participant’s individual space using a non-linear registration (Joshi et al., 2011). The atlas was reduced to 83 nodes after excluding nodes that had missing data. The mean time-course for each of the nodes was calculated and the correlation between each pair of nodes was estimated using a Pearson correlation, resulting in an 83 by 83 connectivity matrix for each infant. Correlations were normalized to *Z* scores using a Fisher transformation.

### 1.5. Adult dataset (HCP 900)

For comparison with the infant data, we used adult data from the Human Connectome Project (HCP) 900 subjects release (Van Essen et al., 2013). The adult MRI (HCP 900) data acquisition details can be found in Ugurbil et al., (2013). For the adult resting-state data (HCP), the resting-state fMRI protocol has been previously described in detail (Glasser et al., 2013; Smith et al., 2013, Van Essen et al., 2013). Resting-state data was acquired on two days; for the current study Day 1 (left-to-right phase encoding) resting-state data was considered Session 1 and Day 2 resting-state data (left-to-right phase encoding) was considered Session 2 as described in Horien et al. (2018).

We obtained the minimally processed HCP data (Glasser et al., 2013). Data were then further preprocessed and connectivity matrices were calculated using a previously described analysis pipeline (Finn et al., 2015, 2017; Shen et al., 2017), except a different atlas was used. To match the infant connectivity matrices, connectivity matrices in the adult were calculated using the 95-node infant atlas (Scheinost-95) registered into adult MNI space rather than the Shen-268 atlas. Of the full HCP 900 sample, 835 individuals had complete resting-state scan data for ‘LR’ phase encoding on both days (Session 1 and Session 2). To be able to directly compare the connectivity matrices between the infants and adults, the infant-specific parcellation was registered into adult MNI space and connectivity matrices were calculated based upon this parcellation. This resulted in 83 by 83 connectivity matrices for the adult data. Participants were excluded from the analysis if their average frame-to-frame motion was greater than 0.15 mm. Thus 81 participants were excluded from the adult sample, resulting in 754 participants for the HCP 900 analyses.

### 1.6. Connectome-based ID via Pearson correlation

The connectome-based ID procedure followed previous studies using this method (Finn et al., 2015; Horien et al., 2019) using scripts obtained from NITRC (https://www.nitrc.org/projects/bioimagesuite). This procedure consisted of creating a database of all connectivity matrices for the dataset. Through an iterative process, each participant from a different session (Session 1 or Session 2) is denoted as the ‘target’. The Pearson correlation between the target and all other matrices in the database is computed. A correct ID occurred when the highest Pearson correlation coefficient is between the target in one session and the same in the second session. This process is repeated until each participant served as the target once and is repeated for all participants, sessions, and database-target combinations. The connectome-based ID procedure resulted in two ID rates (ID rates), reflecting the two possible configurations obtained by exchanging the roles of target and database session. ID rate is computed as the percentage of participants whose identity was correctly predicted out of the total number of participants. We examined if ID rates were above chance-level, permutation testing was used to generate a null distribution such that participant identities were shuffled at random and the ID rate was calculated with the random labels. The ID rates from using the correct labels were compared to the null distribution to test for significance. The ID procedure was conducted between all sessions and all scan types (resting-state and native language task). For between-session ID rates, Session 1 was data collected at the first time point (1–2 months old, resting-state or native language task) and Session 2 was data collected from the second time point (9–10 months old, resting-state or native language task). For within-session ID rates, either resting state data was selected as Session 1 and the native language task was selected as Session 2 or vice versa.

### 1.7. Connectome-based ID via geodesic distance

Using geodesic distance, rather than Pearson correlation, as the distance metric in the ID procedure in adults has been shown to achieve over 95% accuracy with resting-state data (Venkatesh, Jaja, & Pessoa, 2020). Geodesic distance is a distance metric that is non-Euclidean and considers the manifold on which the data lies. The Geodesic Distance ID procedure follows the same basic structure as the Pearson correlation-based ID procedure using the 1-Nearest Neighbor except that geodesic distance is computed as the distance measure rather than Pearson correlation. As with the Pearson-based ID procedure, the geodesic distance ID procedure was conducted between all sessions and all scan types (resting-state versus native language task). The geodesic distance ID procedure was implemented using code (https://github.com/makto-toruk/FC_geodesic) from the original geodesic distance ID study (Venkatesh et al., 2020).

### 1.8. Differential power analysis

For each of the ID rates that reached statistical significance via permutation testing we computed the differential power (DP) as described elsewhere (Finn et al., 2015). DP is an estimate for each edge of the likelihood that within-participant similarity is higher than between-subject similarity. DP is calculated as the product of the z-scored edge values from Session 1 and Session 2 from the same participant is compared to the product of the same edge value from Session and Session 2 from unmatched participants (Finn et al., 2015). If the within-participant product is higher than between-participant product across all the participants in the sample, this edge contributes highly to ID (indicated by a high DP value). DP was calculated for the infant data in which ID rates were significant and calculated for the adult (HCP 900) data.

### 1.9. Test–retest reliability analysis

To examine the role of edge-level test–retest reliability in connectome-based ID in the first year of life, we estimated ICC in Matlab using the Multifactor ICC toolbox (https://github.com/SNeuroble/Multifactor_ICC; (Noble et al., 2017)). The Generalizability Theory (G Theory) framework adopted by this tool has been used to estimate test–retest reliability of functional connectivity in a number of studies (Forsyth et al., 2014; Gee et al., 2015; Noble et al., 2017). We calculated the “absolute reliability” form of the ICC (called the ICC in G Theory), which reflects the amount of variance due to the object of measurement (here, participants) relative to error sources (here, session, interactions between participant and session, and residual error; cf. (Shrout & Fleiss, 1979). Specifically, the toolbox estimated variance due to each factor (participant and session) in a 2-way ANOVA with interactions using the MATLAB function ‘anovan’. Negative variance components were small in their magnitude and set to 0 (Shavelson, Baxter, & Gao, 1993). Test-retest reliability was summarized across the brain by calculating the mean and standard deviation of ICC across all edges. ICC can be interpreted as <0.4 = poor; 0.4–0.59 = fair; 0.60–0.74 = good; and >0.74 = excellent (Cicchetti & Sparrow, 1981). Means, standard deviations, and ranges for ICC were calculated for all between all scan types (resting-state and native language task) and sessions (Session 1 and Session 2). To compare infant test–retest reliability to adult test-test reliability, we calculated the edge-level ICCs for the HCP data. We also examined the correlation between ICC and DP and used a one-tailed Mantel test with 1000 iterations to estimate significance of this association since we hypothesized a positive correlation between ICC and DP.

## 2. Results

### 2.1. Connectome-based ID

Using the Pearson-based connectome ID procedure, we found support for our hypothesis that within-session ID rates would be highest (between 48.94% and 70.83%). However, ID rates for Session 2 were not substantially higher than Session 1 as hypothesized. Between-session ID rates were very low and mostly nonsignificant with the highest rate being for Task1–Task2 (26.67%). **Table 2** shows each ID rate for each task–rest pair for the Pearson-based ID procedure with the raw connectivity matrices. ID rates based on geodesic distance were not substantially higher than the ID rates based on Pearson correlation (see **Table 3**). However, the geodesic distance-based ID rates follow a consistent pattern as the Pearson-based ID rates in that the highest ID rates are for the within-session IDs. Between-session ID rates remained very low and mostly nonsignificant. ID rates for the adult data was 71.6% and 71.4%, consistent with previous studies examining ID rate in the HCP sample using the resting-state data (Horien et al., 2018). In adult connectome-based ID studies, ID rate improvements have been shown using geodesic distance as the distance measure rather than Pearson correlation (Abbas et al., 2021). While we observed some improvements of the similar magnitude for the within-session ID rates, overall, geodesic distance did not improve between-session ID rates.

**Table 2.**
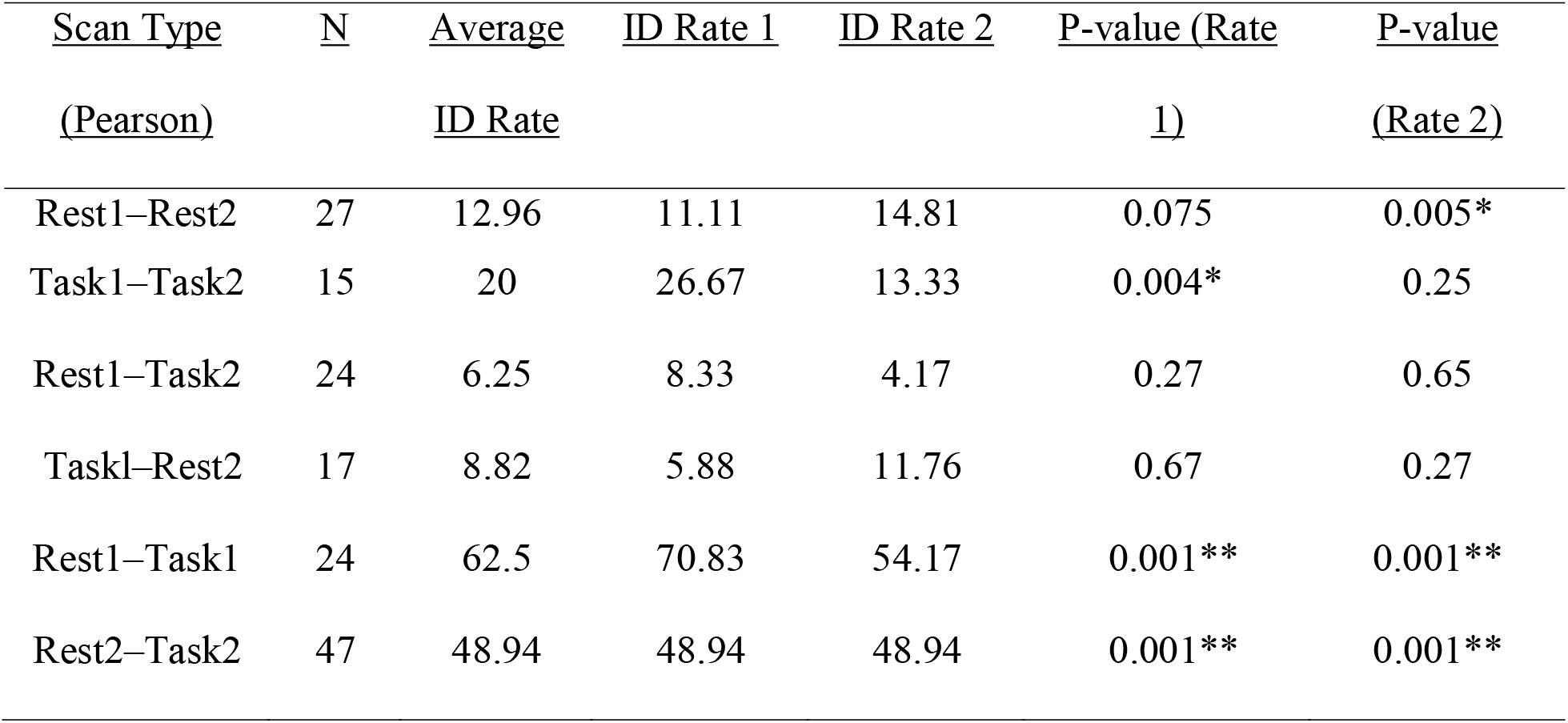
Results of the Pearson connectome-based ID procedure. ID Rate 1 refers to the percentage of correct IDs (a participant at Session 1 was most highly correlated with themselves at Session 2) when using Session 1 as the target and Session 2 as the database, and ID Rate 2 refers to IDs when target and database are reversed.

| Scan Type | N | Average | ID Rate 1 | ID Rate 2 | P-value (Rate | P-value |
| --- | --- | --- | --- | --- | --- | --- |
| (Pearson) |  | ID Rate |  |  | 1) | (Rate 2) |
| Rest1–Rest2 | 27 | 12.96 | 11.11 | 14.81 | 0.075 | 0.005* |
| Task1–Task2 | 15 | 20 | 26.67 | 13.33 | 0.004* | 0.25 |
| Rest1–Task2 | 24 | 6.25 | 8.33 | 4.17 | 0.27 | 0.65 |
| Task1–Rest2 | 17 | 8.82 | 5.88 | 11.76 | 0.67 | 0.27 |
| Rest1–Task1 | 24 | 62.5 | 70.83 | 54.17 | 0.001** | 0.001** |
| Rest2–Task2 | 47 | 48.94 | 48.94 | 48.94 | 0.001** | 0.001** |

**Table 3.**
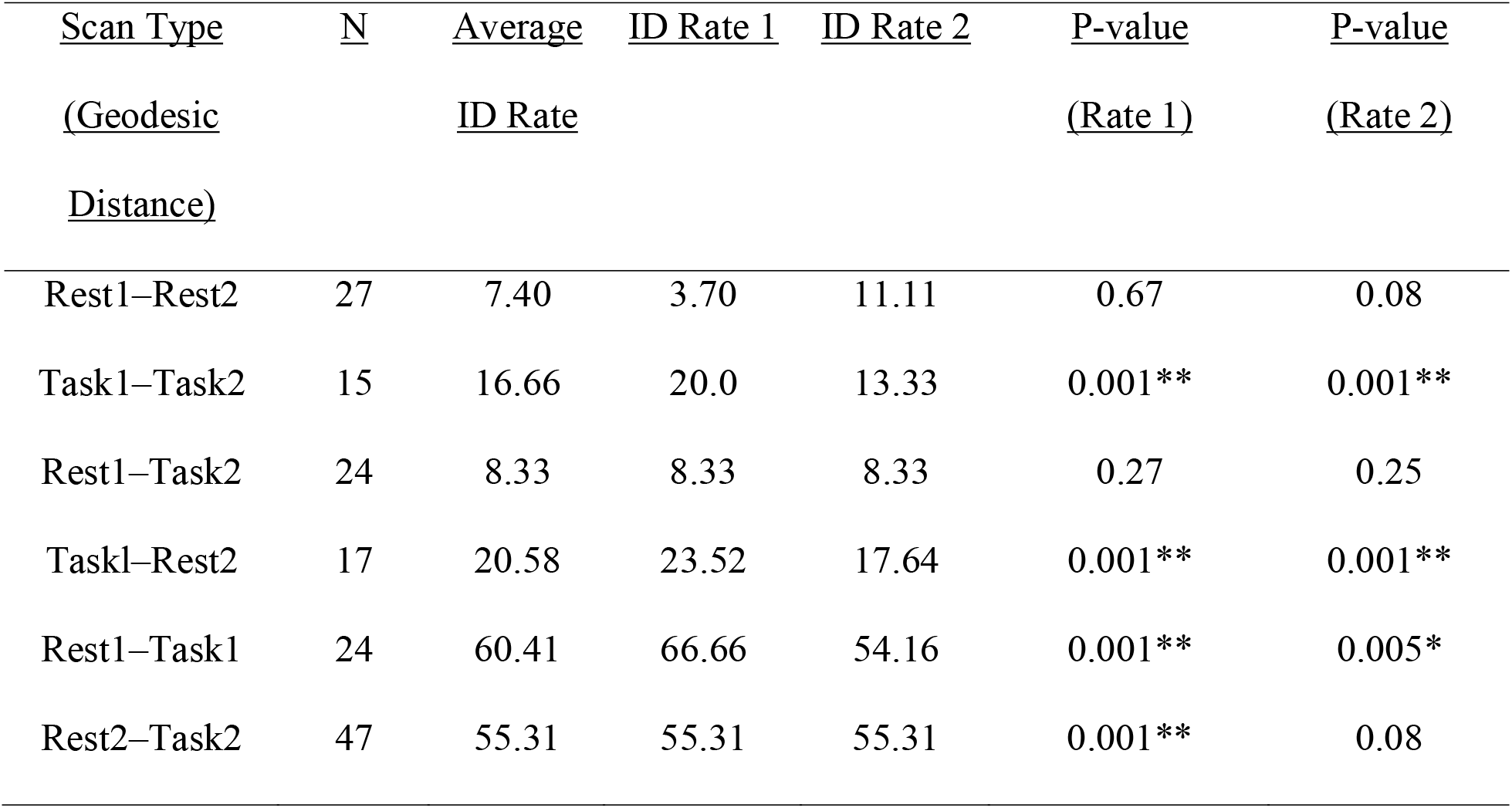
Results of the geodesic distance connectome-based ID procedure. ID Rate 1 refers to the percentage of correct IDs (a participant at Session 1 was most highly correlated with themselves at Session 2) when using Session 1 as the target and Session 2 as the database, and ID Rate 2 refers to IDs when target and database are reversed.

### 2.2. Differential power analysis

DP was calculated for each of the connectome-based ID rates (*p* < 0.05) (see **Figure 1**). The DP analysis focused on the DP for the two within-session ID rates (Rest1–Task1 and Rest2–Task2) for their relatively high ID accuracy. We included a DP analysis of Rest1–Rest2 to examine DP ‘developmentally’ as there was approximately 8 months between the two scans. For comparison, we also calculated DP for the adult data. Similar to previous studies (Finn et al., 2015; Horien et al., 2018; Horien et al., 2019), DP across infant and adult scans all included a widely distributed network of edges including the prefrontal, temporal, and parietal cortices. First, we examined if the DP edges were significantly different between adults and infants in terms of the number of significant prefrontal-to-prefrontal connections (PFC–PFC), prefrontal-to-non-prefrontal (PFC–nonPFC), and non-prefrontal-to-non-prefrontal connections (nonPFC–nonPFC) (see **Table 4** for count totals). Kolmogorov-Smirnov (K-S) tests indicated that DP edge distributions for PFC–PFC connections in Rest1–Task1, Rest1–Rest2, Rest2–Task2 and adult data did not follow a normal distribution (*ps* < 0.001). K-S tests did not reveal any significant differences between adults and infants with respect to the number of DP edges found to be significant (PFC–PFC *D* = 0.01, *p* > 0.05; PFC–nonPFC *D* = 0.02, *p* > 0.05; and nonPFC–nonPFC *D* = 0.05, p > 0.05).”. The same patterns were observed using two a sample K-S test for comparing the adult data with the infant Rest1–Rest2 data (all *p*s > 0.05), and the infant Rest2–Task2 data (all *p*s > 0.05) in terms of number of significant PFC–PFC, PFC–nonPFC, and nonPFC–nonPFC DP edges. These findings suggest that the distributions of DP edges may not meaningfully differ between adults and infants.

**Figure 1.**
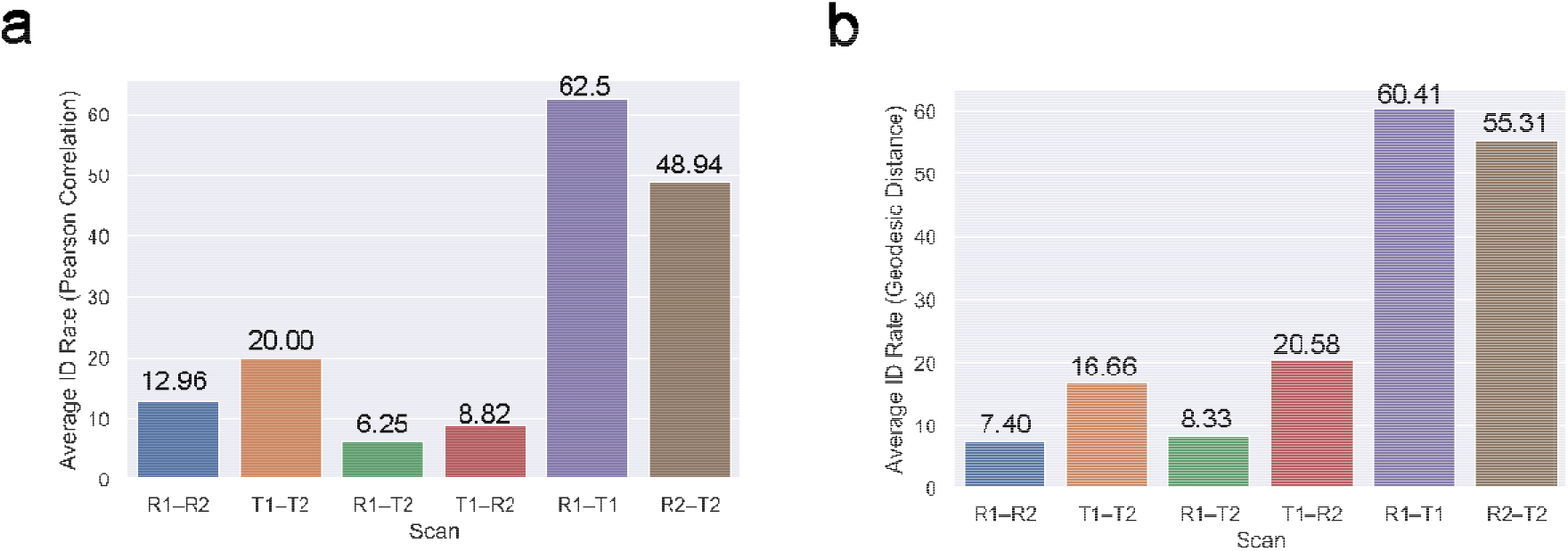
(a) Results of the Pearson connectome-based ID procedure. Bar plots show the average ID rates for each scan pair (R1–R2: Rest1–Rest2, T1–T2: Task1–Task2, R1–T2: (Rest1–Task2), T1–R2: Task1–Rest2, R1–T1: Rest1–Task1, and R2–T2: Rest2–Task2. (b) Results of the geodesic distance connectome-based ID procedure.

**Table 4.** Number of edges with significant differential power (DP; *p* < 0.05) for each connectome-based ID rate.

| <u>Scan</u> | <u>PFC– PFC</u> | <u>PFC–nonPFC</u> | <u>nonPFC–nonPFC</u> |
| --- | --- | --- | --- |
| Infant Rest1–Task1 | 8 | 56 | 81 |
| Infant Rest1–Rest2 | 6 | 52 | 91 |
| Infant Rest2–Task2 | 12 | 65 | 84 |
| Adult Rest1–Rest2 | 9 | 62 | 80 |

Second, we examined if there was a developmental shift of the degree distribution towards higher degree nodes in the prefrontal cortex with the highest being the adult DP data. We calculated the degree for each of the DP matrices and examined if degree distributions were significantly different between adults and infants. A two sample K-S test indicated that there was no significant difference between the degree distributions, across the brain, for the adult data and Rest1–Task1 (*D* = 0.03, *p* < 0.99), adult data and Rest1–Rest2 (*D* = 0.04, *p* < 0.99), and adult data and Rest2–Task2 (*D* = 0.02, *p* < 0.99). Thus, these findings suggest that the degree distributions of DP edges may also not meaningfully differ between adults and infants.

Finally, we examined if the low ID rates in infants may be due to differences in the underlying functional connectivity across the frontoparietal network (as this network has a protracted developmental trajectory (Fair et al., 2007; Gao, Alcauter, Smith, et al., 2015; Peters, Van Duijvenvoorde, Koolschijn, & Crone, 2016) and contributes highly to adult ID rates (Finn et al., 2015; Jalbrzikowski et al., 2019)). For the infant (Rest1–Task1, Rest1–Rest2, and Rest2–Task2) and adult data, we calculated the average functional connectivity across the frontoparietal network (8 nodes) and examined differences between adults’ and infants’ functional connectivity. Using a permutation with 1000 iterations, adult frontoparietal functional connectivity (averaged across Rest1 and Rest) was significantly different than infant frontoparietal functional connectivity (averaged across sessions) for Rest1–Task1 (*t* = −0.028, *p* < 0.001), Rest1–Rest2 (*t* = −0.022, *p* < 0.001), and Rest2–Task2 (*t* = −0.017, *p* < 0.001). There were no significant differences in frontoparietal functional connectivity among the infant data (*p*s > 0.05). The mean frontoparietal functional connectivity for the adults was −0.03 therefore the negative observed differences between the adults and infants (mean for Rest1–Task1 = −0.0013, mean for Rest1–Rest2 = −0.0080, mean for Rest2Task2 = −0.0128) suggest a developmental shift towards more ‘negative’ functional connectivity across the frontoparietal network (with Rest2–Task2, when infants are oldest, showing the least difference from the adult data). Overall, these findings suggest that differences in the underlying functional connectivity may be driving successful ID.

### 2.3. Analysis of successful IDs

For the connectome-based ID rates that were significant: Rest1–Task1, Rest2–Task2, Rest1–Rest2 (Rate 2), Task1–Task2 (Rate 1), we conducted a follow-up analysis of whether successful IDs were influenced by risk status. For Rest1–Task1, successful ID was not found to vary according to infant risk status for ASD (*p* < 0.10, chi square test) or infant sex (*p* < 0.20, chi square test) for Rate 1. For Rest1–Task1 Rate 2, Rest2–Task2 (both rates), Rest1–Rest2 (Rate 2), Task1–Task2 (Rate 1), successful ID did not vary according to infant risk status for ASD or infant sex (all *p*s > 0.05).

### 2.4. Test–retest reliability

We also examined edge-level ICCs for combinations of sessions (summarized in **Table 5**, plotted on the brain in **Figure 2**, and as matrices sorted by lobe in **Figure 3**). For the Pearson-based correlation matrices, within-session mean edge-level ICCs were consistently higher than between-session mean edge-level ICCs. For both types of connectivity matrices, between-session mean edge-level ICCs were consistently low (mean ~0.08). Within-session mean edge-level ICCs were consistently higher for Session 2 when the infants are 9–10 months old versus Session 1 when the infants are 1–2 months old. For both types of connectivity matrices, between-session mean edge-level ICCs were surprisingly not higher during task compared with rest. Overall, across all of the rest-task and session comparisons, the mean edge-level ICCs were in the poor range. However, for both types of connectivity matrices, maximum values reached to ‘good’ or ‘excellent’ range for ICC. For the adult data, the ICC had a mean of 0.38 ±0.12 and a range of 0.04–0.81.

**Figure 2.**
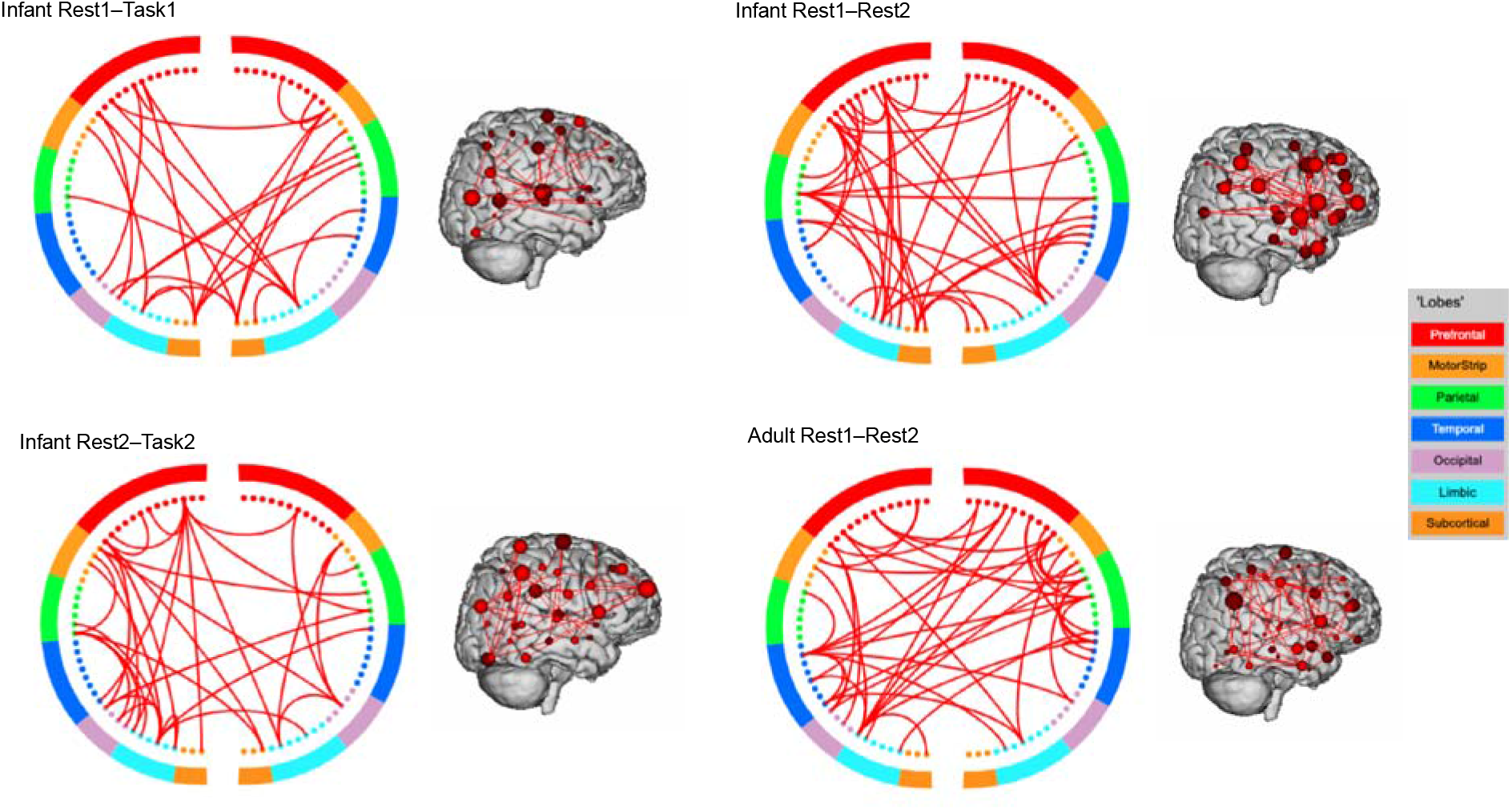
Circle plots showing the significant differential power (DP) edges for each connectome-based ID at *p* < 0.05. For visualization purposes, a degree threshold of 7 was applied. Across infants and adults, patterns are widely distributed involving connections spanning the prefrontal, parietal, temporal, and occipital lobes.

**Figure 3.**
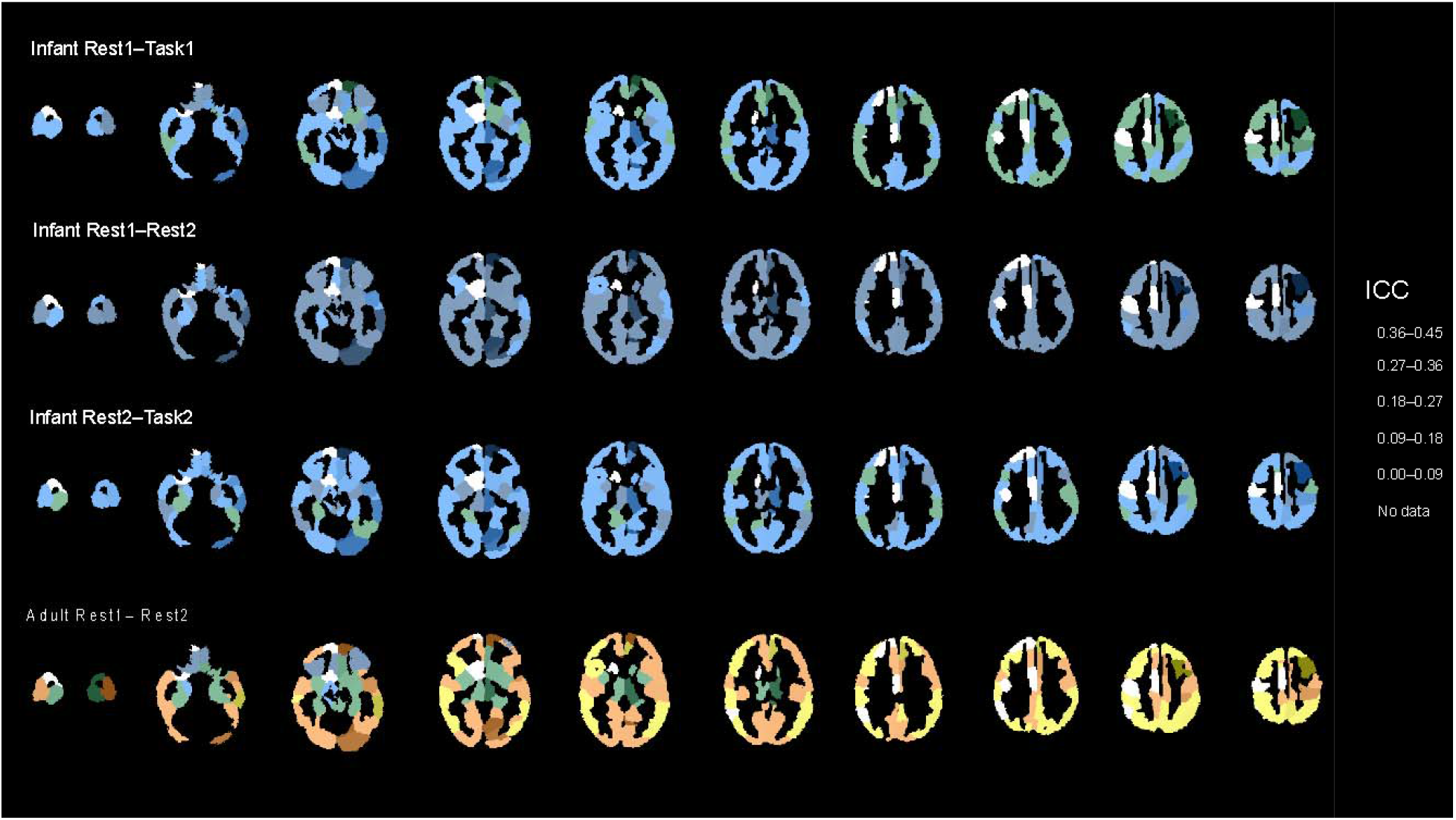
Mean intraclass correlation coefficient (ICC) for each region of the Scheinost-95 parcellation for infants and adults.

**Figure 4.**
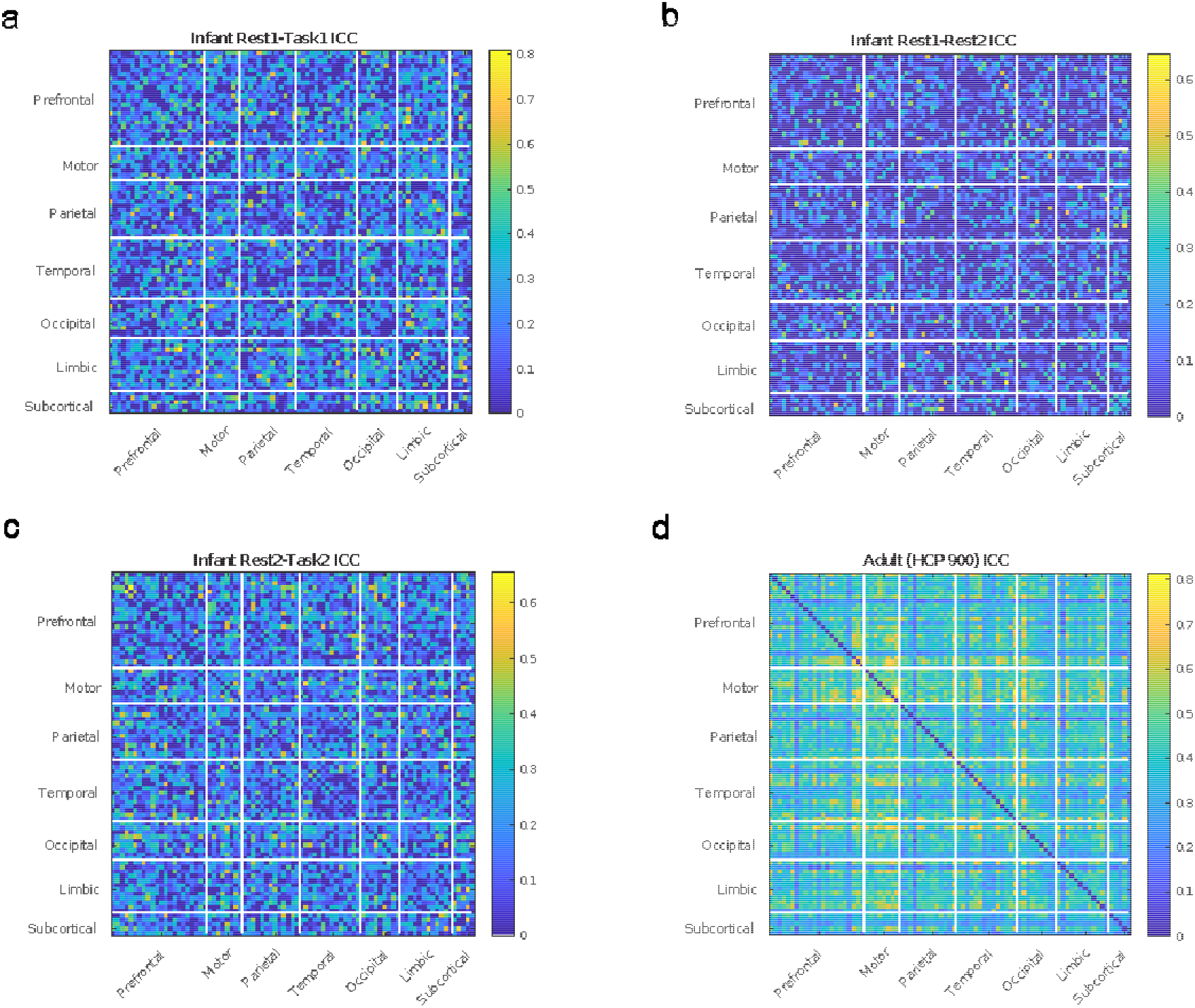
(a) Intraclass correlation coefficients (ICCs) organized by lobe for the infant Rest1–Task1 (b) ICCs organized by lobe for the infant Rest1–Rest2 (c) ICCs organized by lobe for the infant Rest2–Task2 (d) ICCs organized by lobe for the adult data (Rest1–Rest2 for the HCP 900 data).

**Table 5.** Means, standard deviations, and ranges of the edge-level ICC (D-coefficient) for each rest-task and Session combination (Rest1 = resting-state Session 1, Task1 = task Session 1, Rest2 = resting-state Session 2, Task2 = task Session 2) for Pearson correlation connectivity matrices.

| <u>Scan Type</u> | <u>N</u> | <u>ICC (Mean ± SD)</u> | <u>Range</u> |
| --- | --- | --- | --- |
| Rest1–Rest2 | 27 | $0.09 \pm 0.11$ | 0 – 0.64 |
| Task1–Task2 | 15 | $0.14 \pm 0.15$ | 0 – 0.77 |
| Rest1–Task2 | 24 | $0.08 \pm 0.11$ | 0 – 0.55 |
| Task1–Rest2 | 17 | $0.09 \pm 0.13$ | 0 – 0.68 |
| Rest1–Task1 | 24 | $0.19 \pm 0.16$ | 0 – 0.80 |
| Rest2–Task2 | 47 | $0.16 \pm 0.13$ | 0 – 0.65 |

We examined the correlation between ICC matrices for the infants and adults focusing on the two within session ID rates (Rest1–Task1 and Rest2–Task2) and the resting-state between-session ID rate (Rest1–Rest2); the latter was chosen as this session most comparable with the adult data as it was resting-state. To examine the correlations between the matrices, we used a Mantel test (Mantel, 1967) with 1000 iterations. The Mantel test indicated Rest1–Task1 ICC values for the infants were positively correlated with the adult ICC values (*r* = 0.19, *p* < 0.001). A positive correlation was also found between the adult ICC values and infant Rest2–Task2 ICC values (*r* = 0.18, *p* < 0.001). However, there was not a significant association between the adult ICC values and the infant Rest1–Rest2 ICC values (*p* < 0.21).

We examined if there was a significant difference between the ICC values for the adults and infants. Using a permutation test with 1000 iterations, we found that the ICCs for the adult data were significantly greater than the infant Rest1–Task1 data (*t* = 0.17, *p* < 0.001), infant Rest1–Rest2 (*t* = 0.28, *p* < 0.001), infant Rest2–Task2 (*t* = 0.20, *p* < 0.001). The ICCs for the infant Rest1–Rest2 data were lower than the infant Rest2–Task2 (*t* = 0.07, *p* < 0.001) and infant Rest1–Task1 data (*t* = −0.10, *p* < 0.001). However, there were no significant differences between the Rest1–Task1 and Rest2–Task2 data (*p* < 0.99). Together, these results suggest that, while the infant connectome exhibits lower ICC than the adult connectome, similar edges have higher ICC values in infants and adults.

### 2.5. Associations between test–retest reliability and differential power

We estimated the association between edge-level test–retest reliability (ICC) and DP using the Mantel statistic. This enabled us to evaluate the extent to which edges that were the most reliable were the same edges that contributed highly to a successful ID. In all cases, the correlations between the DP and ICC were small (*r*s < 0.04).”

## 3. Discussion

In contrast to prior work in adulthood and adolescence, we found that the infant functional connectome shows poor identifiability across the first year of life. As connectome-based ID rate has been used to quantify the uniqueness and stability of the functional connectome, the results suggest that infants do not yet possess the uniqueness and stability they will have later in development. Within-session ID rates were consistently the highest (48.94–70.83%), with lower ID rates for the between-session analyses (between 14.81–26.67%). As it may be related to ID rate, we also examined edge-level test–retest reliability and found consistent results in which mean ICCs for within-session functional connectome edges were higher (0.16–0.19) while between-session functional connectome edges were consistently lower (0.09–0.14). Further, we found that functional connectome edges in infancy have lower ICCs than adults, however; similar edges have high ICC for infants and adults. These findings have implications for longitudinal studies using fMRI in the first year of life and suggest poor test–retest reliability. The findings show low stability and uniqueness of the functional connectome in infancy and suggest this may be due to the rapid and unparalleled brain development occurring this developmental period. One may expect that low connectome-based ID rates across infancy may be due to poorer test–retest reliability (i.e., less reliable data in infants). However, we did not find support for a correlation between edge-level ICC and DP for the infants. This finding suggests that high DP edges for the infants are not simply edges that have greater test–retest reliability.

Since the first connectome-based ID study was published (Finn et al., 2015), there has been increasing interest in quantifying the uniqueness and stability of individual variability in the functional connectome developmentally. Two studies have shown high ID rates in adolescence (Horien et al., 2019; Jalbrzikowski et al., 2019) and one study found connectome-based ID rates increased between ages 8 to 22 with large increase during puberty (14.5 years old) (Kaufmann et al., 2017). Of particular relevance, both studies in adolescence found that for adolescence, the functional connectome had high ID rates across periods of approximately a year (Horien et al., 2019; Jalbrzikowski et al., 2019). This finding demonstrates high ID rates across a similar range of time as the current study; suggesting that the higher ID rates observed were due to functional connectomes being more developmentally mature in adolescence. The findings from Kaufmann et al., show that ‘connectome distinctiveness’ continues to develop across adolescence. However, it was unclear if high ID rates were achievable from a large age range of 0–8 years old. Studies have quantified rates and development of both the structural and functional development of the brain and highlighted the first year of life as both rapid and unparalleled (Gao et al., 2017; Holland et al., 2014). The results of the current study suggest that for both resting-state and task-based data, within-session connectome-based ID rates tend to be low (below 65%). This pattern suggests that within a session, an infant’s resting-state functional connectome is more highly correlated with its task functional connectome (and vice versa) for about half of the sample. This suggests that even within a session, an infant’s resting-state (or task) connectome is being misidentified and has a higher correlation with another infant’s connectome. For between-session ID rates, greater than 70% of the sample was being misidentified (higher correlation with another individual) for both resting-state and task data. These findings that for the majority of infants in the study, the uniqueness and stability of the functional connectome has not been fully established. In post analysis, we examined if two factors (sex or risk status) were associated with greater ID rates. However, we did not find evidence to support either impacting ID rate.

We did not observe improvements in within-session ID rates (actually lower ID rates) when geodesic distance was used as the distance measure of functional connectivity instead of Pearson correlation (Abbas et al., 2021). The geodesic distance method captures the underlying non-Euclidean geometry of connectivity matrices (Abbas et al., 2021), so perhaps this structure in the connectivity matrix may not be developed enough in infancy (or too noisy) to benefit ID rates. Across the Pearson and geodesic distance ID rates, between-session ID rates were consistently low (below 30%) with within-session ID rates between 40–60%. Future studies will be needed to examine if methods beyond the Pearson and geodesic distance-based ID procedures can achieve high ID rates across the first year of life. We also examined if ID rate varied due to infant sex or ASD risk status. We did not have find evidence for sex-specific effects for ID or that ID rate differed between high risk and low risk infants. However, as discussed further below, frontoparietal connectivity was found to shift towards more ‘adult-like’ patterns in older infants. This suggests individual differences in ID success may be due to the maturity of frontoparietal connections. This hypothesis will need to be tested in future studies in which frontoparietal connectivity and ID rates can be examined longitudinally across infancy.

As connectome-based ID is a measure of multivariate test–retest reliability, we calculated the ICC of the edges of functional connectomes across scan (resting-state versus task) and age (1 month and 9 months). Whether between-session or within-session, mean edge ICC values were in the poor range for the infant data. As a comparison, mean edge ICC values were still in the poor range (0.37) but close the threshold of being considered fair (0.40) (Cicchetti & Sparrow, 1981) and similar to a meta-analysis of 25 resting-state studies that found an average edge-level ICC of 0.29 (Noble, Scheinost, & Constable, 2019). A recent study found similar results in infants (mean postmenstrual age of 45.14) (Wang et al., 2020), with within-session edge-level ICCs between 0.14–0.18. However, this study did not calculate between-session ICCs. These findings suggest that even within-session, infant edge-level ICCs are extremely low and even lower across the first year of life. These findings have implications for future studies examining the infant functional connectome (especially longitudinally) and should be considered when design infant fMRI studies and interpreting results of studies.

We did not find support of our hypothesis that task-based data would have higher edge ICC’s than resting-state data. This may be due to infants being asleep during both rest and the task. However, both Rest1–Task1 and Rest2–Task2 ICCs for the infants were positively correlated with the adult ICCs. This suggests that using a task for infants may make patterns of reliability more similar to the adults. Further analysis of the edge-level ICCs showed a pattern of adult ICCs being higher than all of the infant ICCs, infant Rest1–Rest2 ICCs being lower than all of the other ICCs (infant and adult), and no significant difference between the infant Rest1–Task1 and Rest2–Task2. The Rest1–Rest2 ICCs are consistent with the pattern that between-session ICCs have lower test–retest reliability, these data show it drop substantially across the first year of life and highlight the extensive development occurring during this period.

Connectome-based ID studies in adolescence and adulthood have consistently found the highest contributing edges (high DP) to successful ID were frontoparietal and medial frontal connections (Finn et al., 2015; Horien et al., 2019; Jalbrzikowski et al., 2019). As these connections are present but immature in infancy, it is unclear if high DP edges in infants would differ from adults. DP connectomes were remarkably similar in terms of connections with edges spanning the brain among infants and between adults and infants. We did not find significant differences in the distribution of edges or the degree distribution of edges between adults and infants. These findings suggest that the difference observed between adult ID and infant ID rate is not due to connections that are typically high DP (frontoparietal and medial frontal) being ‘absent’ in regard to not being developed enough in infancy to contribute to fingerprinting. As these primarily prefrontal connections were not absent, we also tested if the count of DP edges between PFC–PFC, PFC–nonPFC, nonPFC–nonPFC were significantly different between infants and adults. The lack of significant difference confirms that the distribution of edges was not different between infants and adults., We examined if the difference in ID rate between infants and adults may be attributable to higher degree distribution in prefrontal nodes such that adults had more developed ‘hubs’ in prefrontal regions and this helped to achieve higher ID rates.

While the results suggest there was not a significant difference DP distribution or degree distribution of DP, we found that the underlying frontoparietal functional connectivity was significantly different in infants compared to adults. The pattern of results comparing adult to infant frontoparietal network connectivity showed a developmental shift towards more negative functional connectivity. Older infants (Rest2–Task2) had the lowest difference between infants and adults while the youngest infant (Rest1–Task1) had the greatest difference between infants and adults. This pattern towards greater negative frontoparietal functional connectivity has been shown in previous developmental studies; however, these typically focus on cortical–subcortical connectivity changes developmentally and less is known about the development of functional connectivity within the frontoparietal network (Lee & Telzer, 2016; Vendetti & Bunge, 2014). In addition to having consistently high contributions to successful ID rates (Finn et al., 2015; Horien et al., 2019; Jalbrzikowski et al., 2019), the frontoparietal network has been shown to develop rapidly across the first year of life (Gao, Alcauter, Elton, et al., 2015; Gao, Alcauter, Smith, et al., 2015) and continue to development into childhood and adolescence (Barber, Caffo, Pekar, & Mostofsky, 2013; Kipping, Tuan, Fortier, & Qiu, 2017; Kolskår et al., 2018). Therefore, the low ID rates in infants compared to adults may be attributable to the differences in overall functional connectivity across the frontoparietal network.

We would like to note a few limitations. First, all of the scans were conducted during the infant’s natural sleep. It is not currently clear how sleep impacts connectome-based ID rate as all studies to data have been conducted in awake participants. Infant functional connectivity during sleep has been shown to most closely resemble adult functional connectivity during sleep (Mitra et al., 2017). Examining ID rates in sleeping adults may be a way to address this limitation as measuring sleep state for infants during scanning would be a large methodological challenged. Second, it will be critical to examine ID rates across infancy in further samples to ensure these findings are not sample specific. The sample was collected to be comprised of ~50% infants with high genetic risk for ASD. We attempted to mitigate this limitation by demonstrating that there was no significant difference in ID rates between low-risk and high-risk infants. Third, the amount of data per acquired for each infant relatively short (7–8 minutes) compared to 12 minutes of data per run collected for the HCP. As total scan length is important for ID rates (Horien et al., 2018), future studies will be needed to examine the impact of run length on ID rates in infancy. Fourth, the imaging sequences used for the data collection have been improved since the data collection (non-multiband sequences). Further, differences in the conditions between task and rest as well as the imaging parameters being difference may have impacted the results. However, previous work has shown that the spatiotemporal resolution of the scan does not impact ID rates (Horien et al., 2018). Still, it is unclear how these factors may have impacted test–retest reliability for infants.

## 4. Conclusions

To our knowledge, this work is the first to show evidence of the low uniqueness and stability of the pattern of functional connectivity (i.e., “fingerprint”) across infancy. This suggests that individual variability of the functional connectome is high across infancy with even more substantial variability within-session. We found that test–retest reliability (for both within- and between-sessions) is poor for infants, with the poorest ICCs found for within-session infant scans. Overall, these findings suggest that in addition to issues with collecting reliable data in infants, individual variability in the functional connectome is high early in development and that low ID rates across this period may reflect the rapid and expansive brain development occurring during this time.

## Acknowledgements

The current study was supported by National Institutes of Health Grant: T32MH18268 (AJD).

## Data Availability Statement

The data for this study is available at nda.nih.gov (NDARCOL0002026 (Susan Bookheimer)).

## References

Abbas, K., Liu, M., Venkatesh, M., Amico, E., Kaplan, A. D., Ventresca, M., … Goñi, J. (2021). Geodesic distance on optimally regularized functional connectomes uncovers individual fingerprints. Brain Connectivity.

Barber, A. D., Caffo, B. S., Pekar, J. J., & Mostofsky, S. H. (2013). Developmental changes in within-and between-network connectivity between late childhood and adulthood. Neuropsychologia, 51(1), 156–167.

Camilli, G., & Hopkins, K. D. (1978). Applicability of chi-square to 2× 2 contingency tables with small expected cell frequencies. Psychological bulletin, 85(1), 163.

Cicchetti, D. V., & Sparrow, S. A. (1981). Developing criteria for establishing interrater reliability of specific items: applications to assessment of adaptive behavior. American journal of mental deficiency.

Fair, D. A., Dosenbach, N. U., Church, J. A., Cohen, A. L., Brahmbhatt, S., Miezin, F. M., … Schlaggar, B. L. (2007). Development of distinct control networks through segregation and integration. Proceedings of the National Academy of Sciences, 104(33), 13507–13512.

Finn, E. S., Shen, X., Scheinost, D., Rosenberg, M. D., Huang, J., Chun, M. M., … Constable, R. T. (2015). Functional connectome fingerprinting: identifying individuals using patterns of brain connectivity. Nature neuroscience, 18(11), 1664–1671.

Forsyth, J. K., McEwen, S. C., Gee, D. G., Bearden, C. E., Addington, J., Goodyear, B., … Olvet, D. M. (2014). Reliability of functional magnetic resonance imaging activation during working memory in a multi-site study: analysis from the North American Prodrome Longitudinal Study. Neuroimage, 97, 41–52.

Gao, W., Alcauter, S., Elton, A., Hernandez-Castillo, C. R., Smith, J. K., Ramirez, J., & Lin, W. (2015). Functional network development during the first year: relative sequence and socioeconomic correlations. Cerebral cortex, 25(9), 2919–2928.

Gao, W., Alcauter, S., Smith, J. K., Gilmore, J. H., & Lin, W. (2015). Development of human brain cortical network architecture during infancy. Brain Structure and Function, 220(2), 1173–1186.

Gao, W., Lin, W., Grewen, K., & Gilmore, J. H. (2017). Functional connectivity of the infant human brain: plastic and modifiable. The Neuroscientist, 23(2), 169–184.

Gee, D. G., McEwen, S. C., Forsyth, J. K., Haut, K. M., Bearden, C. E., Addington, J., … Cornblatt, B. A. (2015). Reliability of an fMRI paradigm for emotional processing in a multisite longitudinal study. Human brain mapping, 36(7), 2558–2579.

Holland, D., Chang, L., Ernst, T. M., Curran, M., Buchthal, S. D., Alicata, D., … Yamakawa, R. (2014). Structural growth trajectories and rates of change in the first 3 months of infant brain development. JAMA neurology, 71(10), 1266–1274.

Horien, C., Noble, S., Finn, E. S., Shen, X., Scheinost, D., & Constable, R. T. (2018). Considering factors affecting the connectome-based identification process: Comment on Waller et al. Neuroimage, 169, 172–175.

Horien, C., Shen, X., Scheinost, D., & Constable, R. T. (2019). The individual functional connectome is unique and stable over months to years. Neuroimage, 189, 676–687.

Jalbrzikowski, M., Lei, F., Foran, W., Calabro, F., Roeder, K., Devlin, B., & Luna, B. (2019). Cognitive and default mode networks support developmental stability in functional connectome fingerprinting through adolescence. BioRxiv, 812719.

Joshi, A., Scheinost, D., Okuda, H., Belhachemi, D., Murphy, I., Staib, L. H., & Papademetris, X. (2011). Unified framework for development, deployment and robust testing of neuroimaging algorithms. Neuroinformatics, 9(1), 69–84.

Kaufmann, T., Alnæs, D., Doan, N. T., Brandt, C. L., Andreassen, O. A., & Westlye, L. T. (2017). Delayed stabilization and individualization in connectome development are related to psychiatric disorders. Nature neuroscience, 20(4), 513–515.

Kipping, J. A., Tuan, T. A., Fortier, M. V., & Qiu, A. (2017). Asynchronous development of cerebellar, cerebello-cortical, and cortico-cortical functional networks in infancy, childhood, and adulthood. Cerebral cortex, 27(11), 5170–5184.

Kolskår, K. K., Alnæs, D., Kaufmann, T., Richard, G., Sanders, A.-M., Ulrichsen, K. M., … Westlye, L. T. (2018). Key brain network nodes show differential cognitive relevance and developmental trajectories during childhood and adolescence. Eneuro, 5(4).

Lee, T.-H., & Telzer, E. H. (2016). Negative functional coupling between the right frontoparietal and limbic resting state networks predicts increased self-control and later substance use onset in adolescence. Developmental Cognitive Neuroscience, 20, 35–42.

Mantel, N. (1967). The detection of disease clustering and a generalized regression approach. Cancer research, 27(2 Part 1), 209–220.

Mitra, A., Snyder, A. Z., Tagliazucchi, E., Laufs, H., Elison, J., Emerson, R. W., … Dager, S. (2017). Resting-state fMRI in sleeping infants more closely resembles adult sleep than adult wakefulness. PloS one, 12(11), e0188122.

Noble, S., Scheinost, D., & Constable, R. T. (2019). A decade of test-retest reliability of functional connectivity: A systematic review and meta-analysis. Neuroimage, 203, 116157.

Noble, S., Spann, M. N., Tokoglu, F., Shen, X., Constable, R. T., & Scheinost, D. (2017). Influences on the test–retest reliability of functional connectivity MRI and its relationship with behavioral utility. Cerebral cortex, 27(11), 5415–5429.

Peters, S., Van Duijvenvoorde, A. C., Koolschijn, P. C. M., & Crone, E. A. (2016). Longitudinal development of frontoparietal activity during feedback learning: Contributions of age, performance, working memory and cortical thickness. Developmental Cognitive Neuroscience, 19, 211–222.

Scheinost, D., Kwon, S. H., Shen, X., Lacadie, C., Schneider, K. C., Dai, F., … Constable, R. T. (2016). Preterm birth alters neonatal, functional rich club organization. Brain Structure and Function, 221(6), 3211–3222.

Shavelson, R. J., Baxter, G. P., & Gao, X. (1993). Sampling variability of performance assessments. Journal of educational Measurement, 30(3), 215–232.

Shrout, P. E., & Fleiss, J. L. (1979). Intraclass correlations: uses in assessing rater reliability. Psychological bulletin, 86(2), 420.

Vendetti, M. S., & Bunge, S. A. (2014). Evolutionary and developmental changes in the lateral frontoparietal network: a little goes a long way for higher-level cognition. Neuron, 84(5), 906–917.

Venkatesh, M., Jaja, J., & Pessoa, L. (2020). Comparing functional connectivity matrices: A geometry-aware approach applied to participant identification. Neuroimage, 207, 116398.

Wang, Y., Hinds, W., Duarte, C. S., Lee, S., Monk, C., Wall, M., … Mamin, M. G. (2020). Intra-session test-retest reliability of functional connectivity in infants. BioRxiv.

Zhang, H., Shen, D., & Lin, W. (2019). Resting-state functional MRI studies on infant brains: a decade of gap-filling efforts. Neuroimage, 185, 664–684.

